# Evaluation of Deep Learning Strategies for Nucleus Segmentation in Fluorescence Images

**DOI:** 10.1101/335216

**Authors:** Juan C. Caicedo, Jonathan Roth, Allen Goodman, Tim Becker, Kyle W. Karhohs, Matthieu Broisin, Molnar Csaba, Claire McQuin, Shantanu Singh, Fabian Thies, Anne E. Carpenter

**Author notes:** corresponding author, Mailing address: 415 Main Street, Cambridge, MA. 02142.

## Abstract

Identifying nuclei is often a critical first step in analyzing microscopy images of cells, and classical image processing algorithms are most commonly used for this task. Recent developments in deep learning can yield superior accuracy, but typical evaluation metrics for nucleus segmentation do not satisfactorily capture error modes that are relevant in cellular images. We present an evaluation framework to measure accuracy, types of errors, and computational efficiency; and use it to compare deep learning strategies and classical approaches. We publicly release a set of 23,165 manually annotated nuclei and source code to reproduce experiments and run the proposed evaluation methodology. Our evaluation framework shows that deep learning improves accuracy and can reduce the number of biologically relevant errors by half.

## 1. Introduction

Image analysis is a powerful tool in cell biology to collect quantitative measurements in time and space with precision, speed, and sensitivity. From image-based assays to high-content screening (1, 2), microscopy images have led to understanding genetic perturbations, pursuing drug discovery, and phenotyping cells in biomedical applications and cell biology research (3, 4). The most widely used quantitative imaging technique in biological laboratories is fluorescence imaging; with automation it can easily produce terabytes of primary research data (5). Accurate and automated analysis methods are key to successfully quantify relevant biology in such large image collections.

One critical step in quantifying fluorescent images is often the identification of the nucleus of each cell with a DNA stain, and there is a long history of research efforts to designing and improving nuclear and cellular segmentation (6). One of the most commonly used strategies for nucleus segmentation is Otsu’s thresholding method (7) followed by seeded watershed (8, 9), because of its effectiveness, simplicity of use and computational efficiency. Machine learning-based segmentation methods have also been introduced for segmenting cells (10), which typically require annotated examples in the form of segmentation masks or interactive scribbles. Many of these strategies are readily available in various bioimage software packages (11), including open source options such as CellProfiler (12), Ilastik (10), and ImageJ/Fiji (13), facilitating their adoption in routine biological research.

Despite widespread adoption, segmentation tools in biology generally do yield non-trivial amounts of segmentation error. These may silently propagate to downstream analyses, yielding unreliable measures or systemic noise that is difficult to quantify and factor out. There are several causes for segmentation errors. First, existing algorithms have limitations due to the assumptions made in the computational design that do not always hold, such as thresholding methods that assume bimodal intensity distributions, or region growing that expects clearly separable boundaries. Second, the most popular solutions for nucleus segmentation were originally formulated and adopted several decades ago when the biological systems and phenotypes of interest were often simpler; however, as biology pushes the limits of high-throughput cellular and tissue models, natural and subtle variations of biologically meaningful phenotypes are more challenging to segment. Finally, algorithms are usually configured using a few –hopefully representative– images from the experiment, but variations in signal quality and the presence of noise pose challenges to the robustness and reliability of the solution at large scale.

The ideal approach to nucleus segmentation would be a generic, robust and fully automated solution that is as reliable as modern face detection technologies deployed in mobile applications and social networks. The current state of the art in face detection and many other computer vision tasks is based on deep learning (14), which has demonstrated high accuracy, even surpassing human-level performance in certain tasks (15). Several models based on deep learning have already been proposed for cell segmentation in biological applications, most notably U-Net (16) and DeepCell (17), which are based on convolutional neural networks.

In this paper, we present an evaluation framework, including a new metric, to answer the question of how much improvement is obtained when adopting deep learning models for nucleus segmentation. The most commonly-used metric for nucleus/cell segmentation evaluation is the Jaccard index (17–19), which measures pixel-wise overlap between ground truth and segmentations estimated by an algorithm. From a cell biology perspective, pixel-wise overlap alone is not useful to diagnose the errors that actually impact the downstream analysis, such as missing and merged objects (Figure S1). Thus, we recommend the use of metrics that explicitly count correctly segmented objects as true positives and penalize any instance-level error, similar to practice in diagnostic applications (20, 21). These include object-level F1-score and false positives, among others.

We demonstrate the utility of this methodology by evaluating two deep learning methods proposed for cell segmentation and comparing them against classical machine learning and image processing algorithms. The goal of our study is to investigate the potential of deep learning algorithms to improve the accuracy of nucleus segmentation in fluorescence images. Expert biologists on our team hand-annotated more than 20,000 nuclei in an image collection of 200 images of the DNA channel from a large image-based chemical screen, sampled from a diverse set of treatments (22). We apply our evaluation framework to analyze different types of segmentation errors, computational efficiency, and the impact of quantity and quality of training data for creating deep learning models.

Our study is restricted to segmenting the nucleus of cells in fluorescence images, which is different from the more general cell segmentation problem. We also assume the availability of a large dataset of annotated images for training the machine learning models. We note that the deep learning techniques evaluated in our experiments were designed for different settings: general purpose cell segmentation with limited training data. Nevertheless, both deep learning methods (DeepCell and U-Net) showed improved ability for segmenting nuclei and fix errors that are relevant to experimental biology when trained with a large image set.

## 2. Materials

### 2.1 Image Collection

The image set is a high-throughput experiment of chemical perturbations on U2OS cells, comprising 1,600 bioactive compounds (22). The effect of treatments was imaged using the Cell Painting assay (23) which labels cell structures using six stains, including Hoechst for nuclei. From this image collection, we randomly sampled 200 fields of view of the DNA channel, each selected from a different compound. By doing so, phenotypes induced by 200 distinct chemical perturbations were sampled.

The original image collection is part of the Broad Bioimage Benchmark Collection, with accession number BBBC022, and the subset used for this study has been made publicly available with accession number BBBC039 at https://data.broadinstitute.org/bbbc/BBBC039/.

### 2.2 Expert Annotations

Each image in the sampled subset was reviewed and manually annotated by PhD-level expert biologists. Annotators were made to label each single nucleus as a distinguishable object, even if nuclei happen to be clumped together or appear to be touching each other. Nuclei of all sizes and shapes were included as our goal was to densely annotate every single nucleus that can be recognized in the sampled images, regardless of its phenotype. In this way, a wide variety of phenotypes was covered, including micronuclei, toroid nuclei, fragmented nuclei, round nuclei, and elongated nuclei, among others (22).

Creating a resource of manually annotated images is time consuming, and existing tools for annotating natural images are not ideal for microscopy. In order to improve and simplify the image annotation experience for experts in our team, we created a prototype annotation tool to assign single-object masks in images. Our user interface allowed the experts to zoom in and out to double check details, and also presented the annotation masks overlaid on top of the original image using a configurable transparency layer. Importantly, our annotation tool was based on assisted segmentation based on superpixels, which are computed on intensity features to facilitate user interactions.

Our prototype tool was useful to collect nucleus annotations for this research; however, significant development is needed to improve it. Alternative methods for collecting image annotations now exist, such as the Quanti.us system (24) for distributing manual annotations to non-expert workers on the internet. This may enable the annotation process for new projects to be scaled up quickly to many more images, and, according to their findings, reaching similar precision to experts when multiple workers provide independent annotation replicates.

## 3. Methods

Identifying nuclei in an image is best framed as an “instance segmentation” problem (25), where the challenge is to find distinct regions corresponding to a single class of objects: the nucleus. Semantic segmentation (26), which splits an image to regions of various classes without requiring objects to be separated, is not helpful for nucleus segmentation because there is only one class, and touching nuclei would not be distinguished from each other. Both of the deep learning strategies evaluated in this paper are cases of instance segmentation that formulate nucleus segmentation as a boundary detection problem.

The boundary detection problem consists of identifying three different types of pixels in an image of nuclei: 1) background, 2) interior of nuclei, and 3) boundaries of nuclei. This formulation simplifies the problem of instance segmentation into a three-class, pixel-wise classification problem (Figure 1), which can be understood as a semantic segmentation solution to identify the structural elements of the image. The critical class to separate single objects is the boundary class: failure to classify boundaries correctly will result in merged or split objects. The actual segmentation masks are obtained from the interior class, which covers the regions where objects are located. This requires additional post-processing steps over classification probability maps to recover individual instances (Suppl. B). Note that while we pose this as a pixel-wise classification problem of boundaries, we evaluate the performance on the success of identifying entire objects.

**Figure 1.**
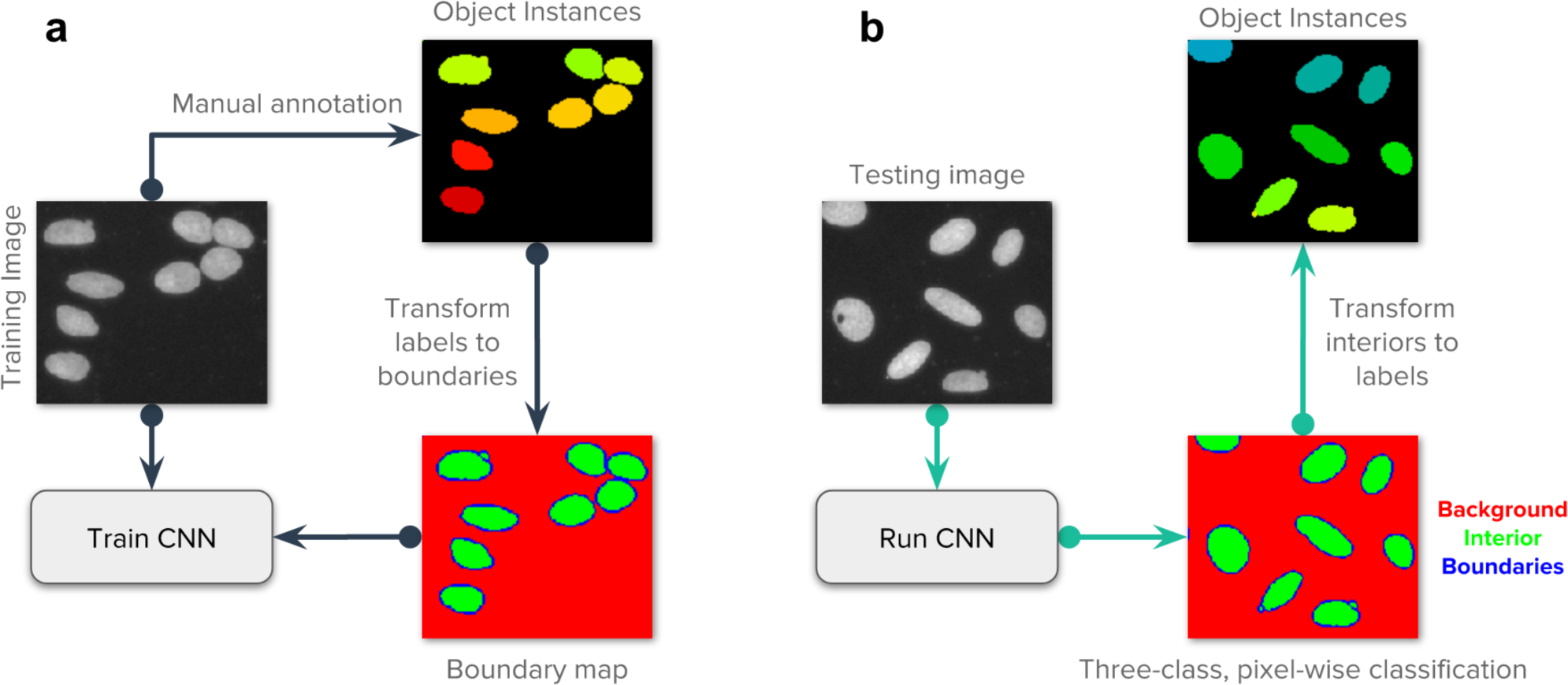
Strategy of the evaluated deep learning approaches. Our main goal is to follow the popular strategy of segmenting each nucleus and micronucleus as a distinct entity, regardless of whether it shares the same cell body with another nucleus. It is generally possible to group nuclei within a single cell using other channels of information in a post-processing step if the assay requires it. a) Images of the DNA channel are manually annotated, labeling each nucleus as a separate object. Then, labeled instances are transformed to masks for background, nucleus interior and boundaries. A convolutional neural network (CNN) is trained using the images and their corresponding masks. b) The trained CNN generates predictions for the three class classification problem. Each pixel belongs to only one of the three categories. In post-processing, the predicted boundary mask is used to identify each individual instance of a nucleus.

The ground truth to solve the boundary detection problem starts with masks individually assigned to each object (Section 2.2) and then transformed to three-class annotations. The boundary annotations are initially obtained from ground truth annotations using a single pixel contour around each nucleus. We then expand this contour with two more pixels, one inside and another outside the boundary to cover natural pixel intensity variations of the input image and also to compensate for errors in manual annotations. We tested with boundaries of different sizes but did not observe any significant benefits of further adjusting this configuration.

Various convolutional neural networks (CNN) can be used for nucleus segmentation, including the Mask-RCNN model (27), which decomposes images into regions first and then predicts object masks. The automatic design of neural networks by searching the space of architectures with an optimization procedure (28) may also be used to create generic nucleus segmentation networks. However, in this work we do not propose novel architectures, instead, we develop an evaluation methodology to identify errors and improve performance of existing models. Specifically, we evaluate two architectures, representing two prominent models designed and optimized for segmenting biological images: DeepCell (17) and U-Net (16). We use the same preprocessing and postprocessing pipeline when evaluating both CNN models (Suppl. B), so differences in performance are explained by architectural choices only.

### 3.1 DeepCell

DeepCell (17) is a framework designed to perform general purpose biological image segmentation using neural networks. Its driving philosophy is to get deep learning operational in biological laboratories using well constructed models that can be trained with small datasets using modest hardware requirements, making them usable in a small data regime. The DeepCell library is open source and features a Docker container with guidelines for training and testing models with new data via Jupyter Notebooks, and more recently, user friendly functionalities to run a deep learning application in the cloud (29).

The DeepCell model evaluated in our work is a CNN that segments images of cells using a patch-based, single pixel classification objective. The network architecture has seven convolutional layers, each equipped with a ReLu nonlinearity (30) and batch normalization (31); three max-pooling layers to progressively reduce the spatial support of feature maps; and two fully connected layers; totalling about 2.5 million trainable parameters per network (a full DeepCell model is an ensemble of 5 networks). This architecture has a receptive field of 61×61 pixels, which is the approximate area needed to cover a single cell (of a diameter up to 40μm, imaged at 20x at a resolution of 0.656μm/pixel), and produces as output a three-class probability distribution for the pixel centered in the patch.

In our evaluation, we use the recommended configuration reported by Van Valen et al. (17), which was demonstrated to be accurate on a variety of cell segmentation tasks, including mammalian cell segmentation and nuclei. Their configuration include training an ensemble of five replicate networks to make predictions in images. The final segmentation mask is the average of the outputs produced by each individual network. The ensemble increases processing time and the number of trainable parameters, but can also improve segmentation accuracy. The settings of the DeepCell system that we used in our experiments can be reproduced using the following Docker container: https://hub.docker.com/r/jccaicedo/deepcell/.

### 3.2 U-Net

The U-Net architecture (16) resembles an autoencoder (32) with two main sub-structures: 1) an encoder, which takes an input image and reduces its spatial resolution through multiple convolutional layers to create a representation encoding. 2) A decoder, which takes the representation encoding and increases spatial resolution back to produce a reconstructed image as output. The U-Net introduces two innovations to this architecture: First, the objective function is set to reconstruct a segmentation mask using a classification loss; and second, the convolutional layers of the encoder are connected to the corresponding layers of the same resolution in the decoder using skip connections.

The U-Net evaluated in our work consists of eight convolutional layers and three max pooling layers in the encoder branch, and eight equivalent convolutional layers with upscaling layers in the decoder branch. All convolutional layers are followed by a batch normalization layer, and the skip connections copy the feature maps from the encoder to the decoder. The receptive field of the U-Net is set to 256 × 256 pixels during training, which can cover a large group of cells at the same time. This architecture has a total of 7.7 million trainable parameters.

We adapted the objective function as a weighted classification loss, giving 10 times more importance to the boundary class. We apply basic data augmentation during training, including random cropping, flips, 90 degree rotations, and illumination variations. Also, we apply additional data augmentation using elastic deformations, as discussed by the authors (16). The training parameters for this network were tuned using the training and validation sets, and the final model is applied to the test set to report performance. The source code of our U-Net implementation can be found in https://github.com/carpenterlab/2019_caicedo_cytometryA, with an optional CellProfiler 3.0 plugin of this nucleus-specific model (33). Also, a U-Net plugin was independently developed for ImageJ for running generic cell segmentation and quantification tasks (34).

### 3.3 Evaluation metrics

Measuring the performance of cell segmentation has been generally approached as measuring the difference between two segmentation masks: a reference mask with ground truth objects representing the true segmentation, vs. the predicted/estimated segmentation mask. These metrics include root-mean-square deviation (35), Jaccard index (17), and bivariate similarity index (18), among others. However, these metrics focus on evaluating pixel-wise segmentation accuracy only, and fail to quantify object-level errors explicitly (such as missed or merged objects).

In our evaluation, we adopt an object-based accuracy metric that uses a measure of area coverage to identify correctly segmented nuclei. Intuitively, the metric counts the number of single objects that have been correctly separated from the rest using a minimum area coverage threshold. The metric relies on the computation of intersection-over-union between ground truth objects T and estimated objects E:

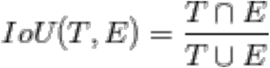

Consider *n* true objects and *m* estimated objects in an image. A matrix *C_n×m_* is computed with all IoU scores between true objects and estimated objects to identify the best pairing. This is a very sparse matrix because only a few pairs share enough common area to score a non-zero IoU value.

To complete the assignment, a threshold greater than 0.5 IoU is applied to the matrix to identify segmentation matches. In our evaluation, we do not accept overlapping objects, i.e., one pixel belongs only to a single nucleus. Thus, a threshold greater than 0.5 ensures that for each nucleus in the ground truth there is no more than one match in the predictions, and vice versa. We also interpret this threshold as requiring that at least half of the nucleus is covered by the estimated segmentation to call it a true positive. In other words, segmentations smaller than half the target object are unacceptable, raising the bar for practical solutions. This differs from a previous strategy (Singh et al. 2009) that takes a similar approach but does not impose this constraint, potentially allowing for very low object coverage. At a given IoU threshold *t*, the object-based segmentation F1-score is then computed as:

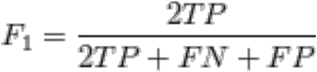

We compute the average F1-score across all images, and then across multiple thresholds, starting at *t* = 0.50 up to *t* = 0.90 with increments △*t* = 0.05. This score summarizes the quality of segmentations by simultaneously looking at the proportion of correctly identified objects as well as the pixel-wise accuracy of their estimated masks.

Our evaluation metric is similar in spirit to other evaluation metrics used in computer vision problems, such as object detection in the PASCAL challenge (36) and instance segmentation in the COCO challenge (25, 37). One important difference of these metrics and ours is that our problem considers a single object category (the nucleus), and therefore, it is more convenient to adopt the F1-score instead of precision.

In our evaluation, we also measure other quality metrics, including the number and type of errors to facilitate performance analysis (38). The following are different types of errors that a segmentation algorithm can make: false negatives (missed objects); merges (under-segmentations), which are identified by several true objects being covered by a single estimated mask; and splits (over-segmentations), which are identified by a single true object being covered by multiple estimated masks. We identify these errors in the matrix *C* of IoU scores using a fixed threshold for evaluation, and keep track of them to understand the difficulties of an algorithm to successfully segment an image.

### 3.4 Baseline Segmentations

#### 3.4.1 Classic Image Processing

We use CellProfiler 3.0 (33) pipelines to create baseline segmentations. CellProfiler was used as a baseline over other tools because it offers great flexibility to configure multi-step image processing pipelines that connect different algorithms for image analysis, and it is widely used in biology labs and high-throughput microscopy facilities. The pipelines are configured and tested by an expert image analyst using images from the training set, and then run in the validation and test set for evaluation. We refer to two CellProfiler pipelines for obtaining baseline segmentations: basic and advanced.

The basic pipeline relies only on the configuration of the module *IdentifyPrimaryObjects*, which is frequently used to identify nuclei. The module combines thresholding techniques with area and shape rules to separate and filter objects of interest. This is the simplest way of segmenting nuclei images when the user does not have extensive experience with image analysis operations, yet it is complete enough to allow them to configure various critical parameters.

The advanced pipeline incorporates other modules for preprocessing the inputs and postprocessing the outputs of the *IdentifyPrimaryObjects* module. In our advanced configuration we included illumination correction, median filters and opening operations, to enhance and suppress features in the input images before applying thresholding. These operations are useful to remove noise and prepare images to the same standard for segmentation using the same configuration. The postprocessing steps include measuring objects to apply additional filters and generate the output masks.

A single pipeline was used for segmenting images in the BBBC039 dataset, while Van Valen’s set required to split the workflow in two different pipelines. We observed large signal variation in Van Valen’s set given that these images come from different experiments and reflect realistic acquisition modes. Two settings were needed for thresholding, the first for normal single mode pixel intensity distributions and another one for bimodal distributions. The latter is applied to cases where subpopulations of nuclei are significantly brighter than the rest, requiring two thresholds. We used a clustering approach to automatically decide which images needed which pipeline. The pipelines used in our experiments are released together with the data and code.

#### 3.4.2 Segmentation using classic machine learning

We used Ilastik (10) to train a supervised machine learning model as an additional benchmark in this evaluation. Ilastik is effective at balancing memory and CPU requirements, allowing users to run segmentations in real time with sparse annotations over the images (scribbles). We loaded the full set of existing annotated training images, and trained a Random Forest classifier with the default parameters. The feature set included intensity, edge, and texture features in 3 different scales. The annotated images used the same three categories used for deep-learning-based segmentation: background, interior and boundaries of nuclei. In addition, we subsampled pixels in the background and interior categories to balance annotations and reduce the impact of redundant pixels.

After the Random Forest model is trained, the validation and test images were loaded into Ilastik for predictions. We obtained the probability maps for each category and applied the same post-processing steps used for deep-learning-based segmentations (Suppl. B).

## 4. Results

### 4.1 Deep learning improves nucleus segmentation accuracy

Overall, we find that deep learning models exhibit higher accuracy than classical segmentation algorithms, both in terms of the number of correctly identified objects, as well as the localization of boundaries of each nucleus (Figure 2). We evaluate these properties using the F1-score (the harmonic average of precision and recall) averaged over increasingly stringent thresholds of overlap between the ground truth and prediction. U-Net and DeepCell obtained higher average F1-scores (Figure S1), yielding 0.898 and 0.858 respectively, versus 0.840 for Random Forests, 0.811 for advanced CellProfiler and 0.790 for the basic CellProfiler pipeline. This improvement is a significant margin when experiments are run at large scale with thousands of images. Deep learning models yield a higher average F1-score across higher thresholds (Figure 2a), indicating that the boundaries of objects are more precisely mapped to the correct contours compared to the other methods.

**Figure 2.**
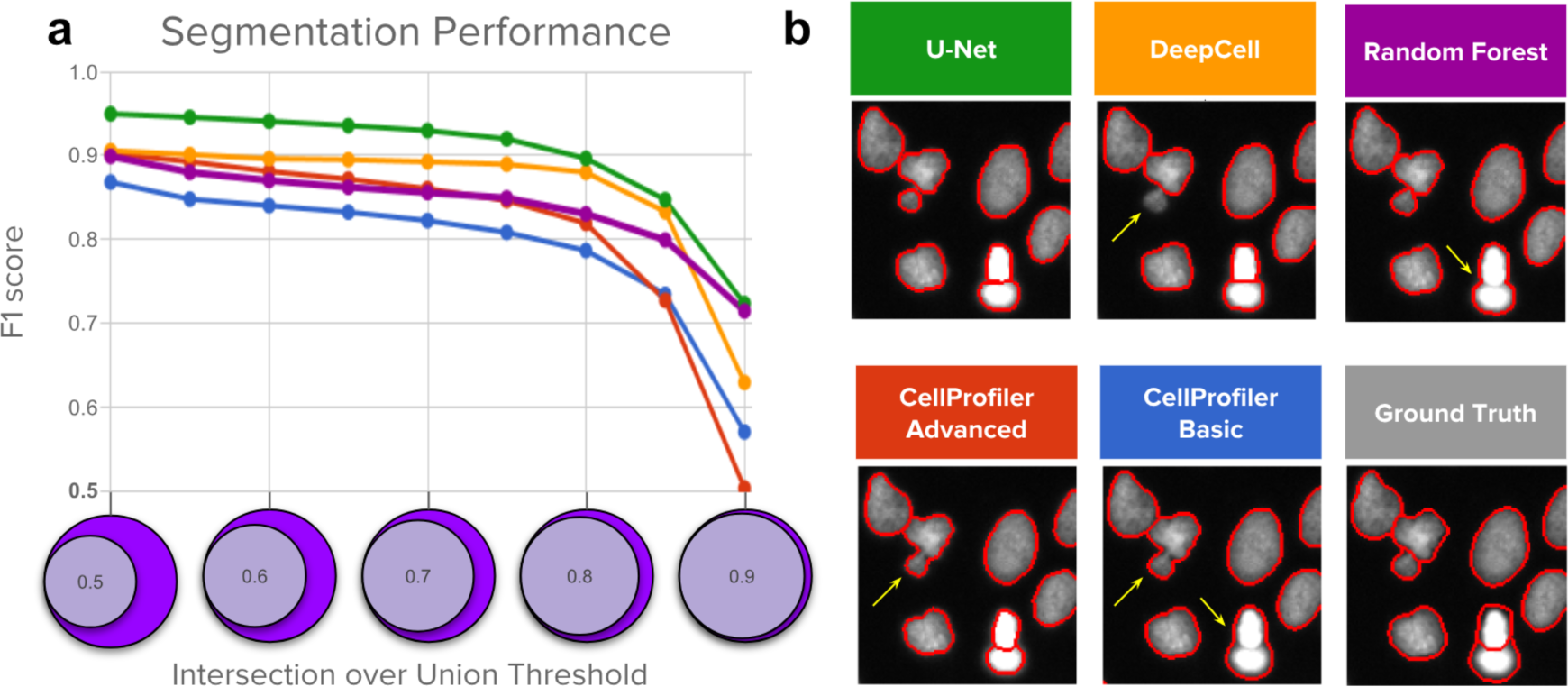
Segmentation performance of five strategies compared against ground-truth expert segmentations. a) Average F1-score vs. nucleus coverage for U-Net (green), DeepCell (yellow), Random Forest (purple), CellProfiler advanced (red), and CellProfiler basic (blue). The y axis is average F1-score (higher is better), which measures the proportion of correctly segmented objects. The x axis represents intersection-over-union (IoU) thresholds as a measurement of how well aligned the ground truth and estimated segmentations must be to count a correctly detected nucleus. Higher thresholds indicate stricter boundary matching. Notice that average F1-scores remain nearly constant up to IoU = 0.80; at higher thresholds, performance decreases sharply, which indicates that the proportion of correctly segmented objects decreases when stricter boundaries are required to count a positive detection. b) Example segmentations obtained with each of the five evaluated methods sampled to illustrate performance differences. Segmentation boundaries are in red, and errors are indicated with yellow arrows.

The most common errors for all methods are merged objects, which occur when the segmentation fails to separate two or more touching nuclei (yellow arrows in Figure 2b). Deep learning strategies tend to reduce this type of error (more in Figure 3c) and provide tighter and smoother segmentation boundaries than those estimated by global Otsu thresholding and declumping, which is at the core of the baseline CellProfiler pipelines for nucleus segmentation.

**Figure 3.**
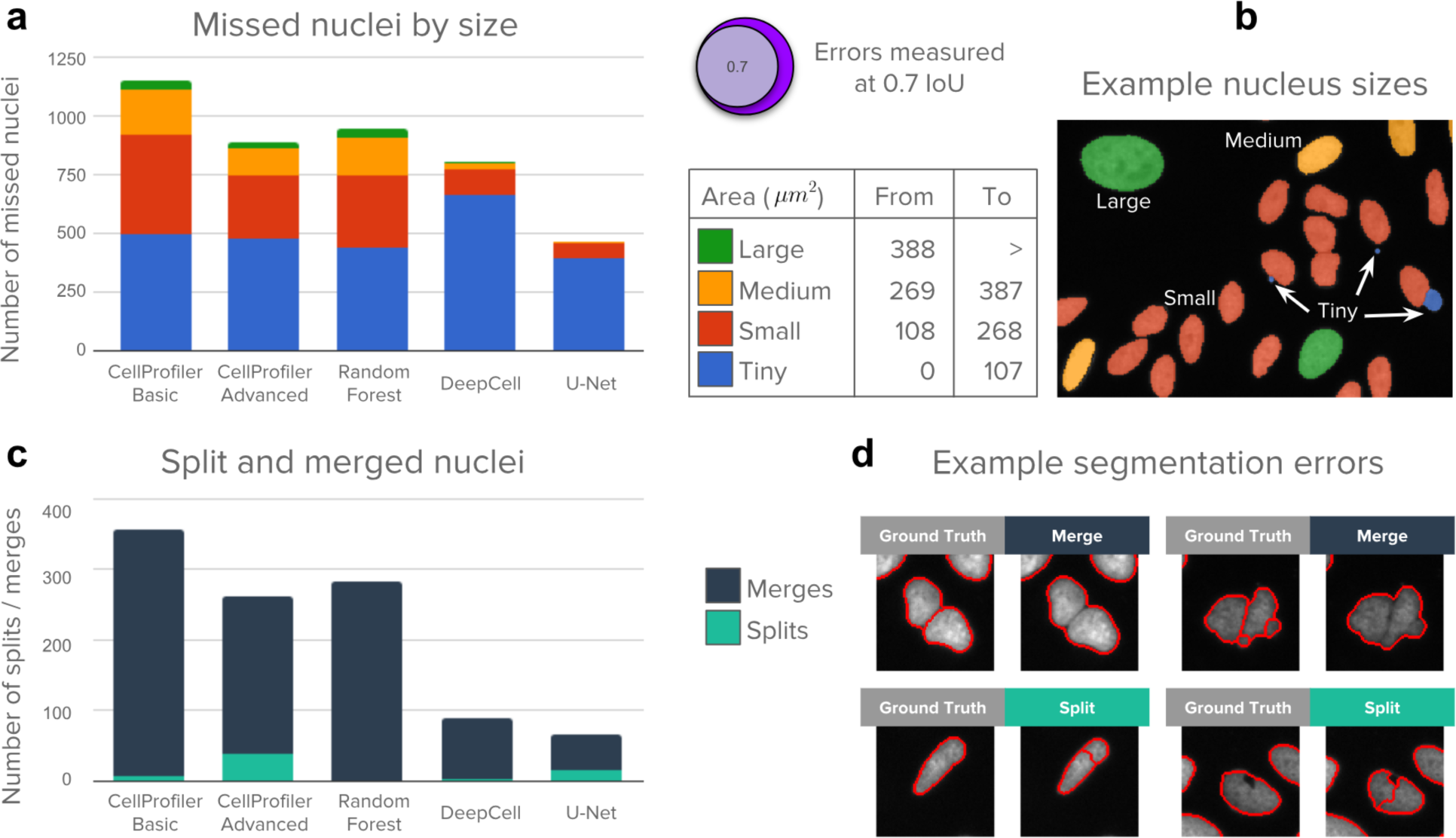
Analysis of segmentation errors (missed, splits, merged objects). The 5,720 nuclei in the test set were used in this analysis. a) Counts of missed nuclei by object size (see table). Missed objects in this analysis were counted using an IoU threshold of 0.7, which offers a good balance between strict nucleus coverage and robustness to noise in ground truth annotations. b) Example image illustrating sizes of nuclei. c) Counts of merged and split nuclei. These errors are identified by masks that cover multiple objects with at least 0.1 IoU. d) Example merges and splits.

Qualitatively, nucleus boundaries predicted by deep learning appear to define objects better than those produced by human annotators using an assistive annotation tool, which can introduce boundary artifacts. Neural nets can learn to provide edges closer to the nuclei with fewer gaps and better-delineated shapes, despite being trained with examples that have such boundary artifacts, showing ability to generalize beyond noise. Overcoming the limitation of assisted annotations is a major strength of this approach because fixing boundary artifacts by hand in the training data is very time consuming. We suspect that the accuracy drop observed in the segmentation performance plot at IoU = 0.85 (Figure 2a) may be partly explained by inaccurate boundaries in ground truth annotations, i.e. improved segmentations may be unfairly scored at high thresholds.

### 4.2 Deep learning excels at correct splitting of adjacent nuclei

Deep learning methods make fewer segmentation mistakes compared to classical pipelines, effectively correcting most of their typical errors (Figure 3a). Here, an error is defined as when a nucleus in the ground truth is missed in an estimated segmentation mask after applying a minimum IoU threshold of 0.7. By this metric, U-Net achieves an error rate of 8.1%, DeepCell 14.0%, Random Forest 16.5%, advanced CellProfiler 15.5% and basic CellProfiler 20.1% (Figure S2). These results are consistent with the evaluation of accuracy performed at multiple IoU thresholds, indicating that deep learning obtains improved performance.

To understand the performance differences among the evaluated methods, we categorized missed objects by size (Figure 3a, b) and segmentation errors by type (merges vs. splits) (Figure 3b, c). An object is missed when the segmentation does not meet the minimum IoU threshold criterion. A merge is counted when one estimated mask is found covering more than one ground truth mask. Similarly, a split is counted when a ground truth mask is being covered by more than one estimated mask. Note that splits and merges are a subset of the total number of errors, and partially overlap with the number of missed objects. That is, some splits and all merges result in one or more missing objects, but not all missing objects are a result of a split or merge.

Deep learning corrects almost all of the errors made by classical pipelines for larger nuclei. We also note that all methods usually fail to capture tiny nuclei correctly (generally, micronuclei, which are readily confounded with debris or artifacts, and represent about 15% of all objects in the test set) (Figure 3a). Interestingly, deep learning tends to accumulate errors for tiny nuclei only, while the CellProfiler pipelines and Random Forests tend to make errors across all sizes (Figure 3b). Tiny nuclei are missed for several reasons, including merging with bigger objects, confounding with debris, or failing to preserve enough object signal for the post-processing routines. Some of these issues can be addressed by developing multi-scale segmentation methods or by increasing the resolution of the input images either optically or computationally.

Both deep learning approaches are effective at recognizing boundaries to separate touching nuclei and correct typical error modes of classical algorithms: merges and splits (Figure 3c and 3d). Split errors produced by the advanced CellProfiler pipeline reveal a trade-off when configuring the parameters of classical algorithms: in order to fix merges we have to accept some more splits. A similar situation happens with U-Net: it has learned to separate clumped nuclei very effectively because the boundary class has 10 times more weight in the loss function, which at the same time forces the network to make some splits to avoid the cost of missing real boundaries (Figure S2).

### 4.3 More training data improves accuracy and reduces errors

We found that training deep learning models with just two images performs already more accurately than an advanced CellProfiler pipeline (Figure 4a). This is consistent with previous experiments conducted in the DeepCell study (17), demonstrating that the architecture of neural networks can be designed and optimized to perform well in the small data regime. Since training a convolutional neural network requires the additional effort of manually annotating example images for learning, limiting the investment of time from expert biologists is valuable.

**Figure 4.**
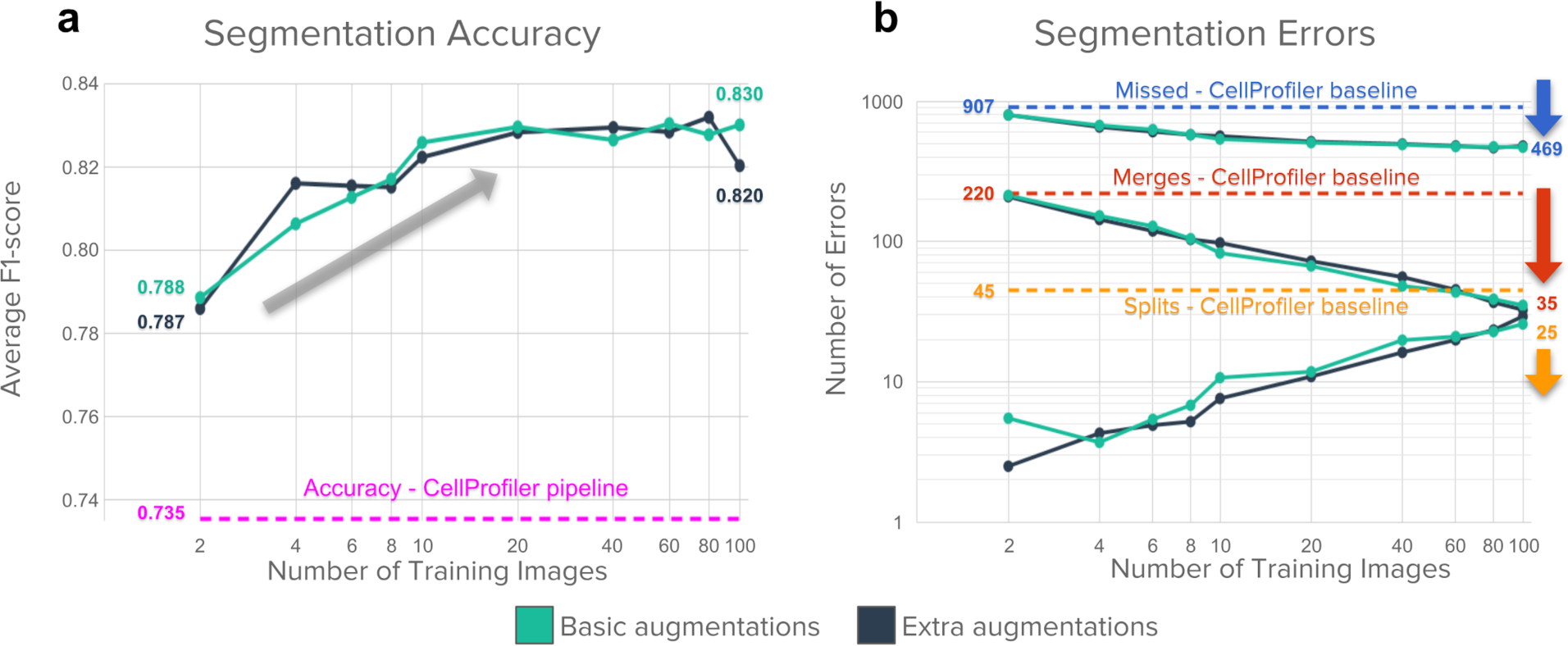
Impact of the number of annotated images used for training a U-Net model. Basic augmentations include flips, 90 degree rotations and random crops. Extra augmentations include the basic plus elastic deformations. a) Accuracy improves as a function of the number of training images, up to a plateau around 20 images (representing roughly 2,000 nuclei). b) Segmentation errors are reduced overall as the number of training images increases, but the impact differs for merges vs. splits. The advanced CellProfiler pipeline is shown as dotted lines throughout. Results are reported using the validation set to prevent over-optimizing models in the test set (holdout). For all experiments, we randomly sampled (with replacement) subsets (n = 2, 4, 6, 8, 10, 20, 40, 60, 80, 100) of images from the training set (n = 100) and repeated 10 times to evaluate performance. Data points in plots are the mean of repetitions. Although the percent overlap between the random samples increases with increasing sample size, and is 100% for n = 100, we nonetheless kept the number of repeats fixed (=10) for consistency. The numbers below each arrow indicate the reduction in number of errors for each category of errors

Data augmentation plays an important role for achieving good generalization results with a small number of images. DeepCell classifies the center pixel of cell-sized patches, creating a large dataset with thousands of examples obtained from each image. Basic augmentations include image rotations, flips, contrast and illumination variations. When combined all together, it results in thousands of training points drawn from an image manifold around the available annotated examples. U-Net follows a similar approach but using larger crops and additional data augmentation based on elastic deformations.

Providing more annotated examples improved segmentation accuracy and reduced the number of errors significantly (Figure 4). Accuracy improves with more data, gaining a few points of performance as more annotated images are used, up to the full 100 images in the training set (Figure 4a). We found little difference in this trend whether using basic data augmentation vs. using extra augmentations based on elastic deformations.

Segmentation errors are reduced significantly with more annotated examples, by roughly half (Figure 4b), but as above, even training with two images produces results better than the advanced CellProfiler baseline. Touching nuclei particularly benefit from more training data, which helps to reduce the number of merge errors. As a model learns to fix difficult merge errors by correctly predicting boundaries between touching objects, a trade-off occurs: some split errors appear in ambiguous regions where no boundaries should be predicted. This effect makes the number of split errors increase with more data, albeit at a slower rate and representing a very small fraction of the total number of errors, while being still fewer than the number of splits made by the advanced CellProfiler pipeline.

### 4.4 Providing a variety of training images improves generalization

Preventing overfitting is an important part of training deep learning models. In our study, we followed the best practices, discussed in detail in the DeepCell work, to mitigate the effect of overfitting: 1) collection of a large annotated dataset for training; 2) the use of pixel normalization and batch normalization to control data and feature variations respectively; 3) the adoption of data augmentation; 4) weight decay to control parameter variations. Besides all these known strategies, we also found that including example images with technical variations in the training set (such as experimental noise or the presence of artifacts) can prevent overfitting, serving as an additional source of regularization.

We found that training with images that exhibit different types of noise produces models that transfer better to other sets (Figure 5a). In contrast, training on an image set that has homogeneous acquisition conditions does not transfer as well to other experiments (Figure 5b). We tested U-Net models trained on one image set and evaluated their performance when transferring to another set. In one case we took images of nuclei from a prior study (“Van Valen’s Set”)(17), representing different experiments and including representative examples of diverse signal qualities, cell lines, and acquisition conditions (Figure 5d). In the other case we used the BBBC039 image collection which exhibits homogeneous signal quality and was acquired under similar technical conditions (Figure 5c).

**Figure 5.**
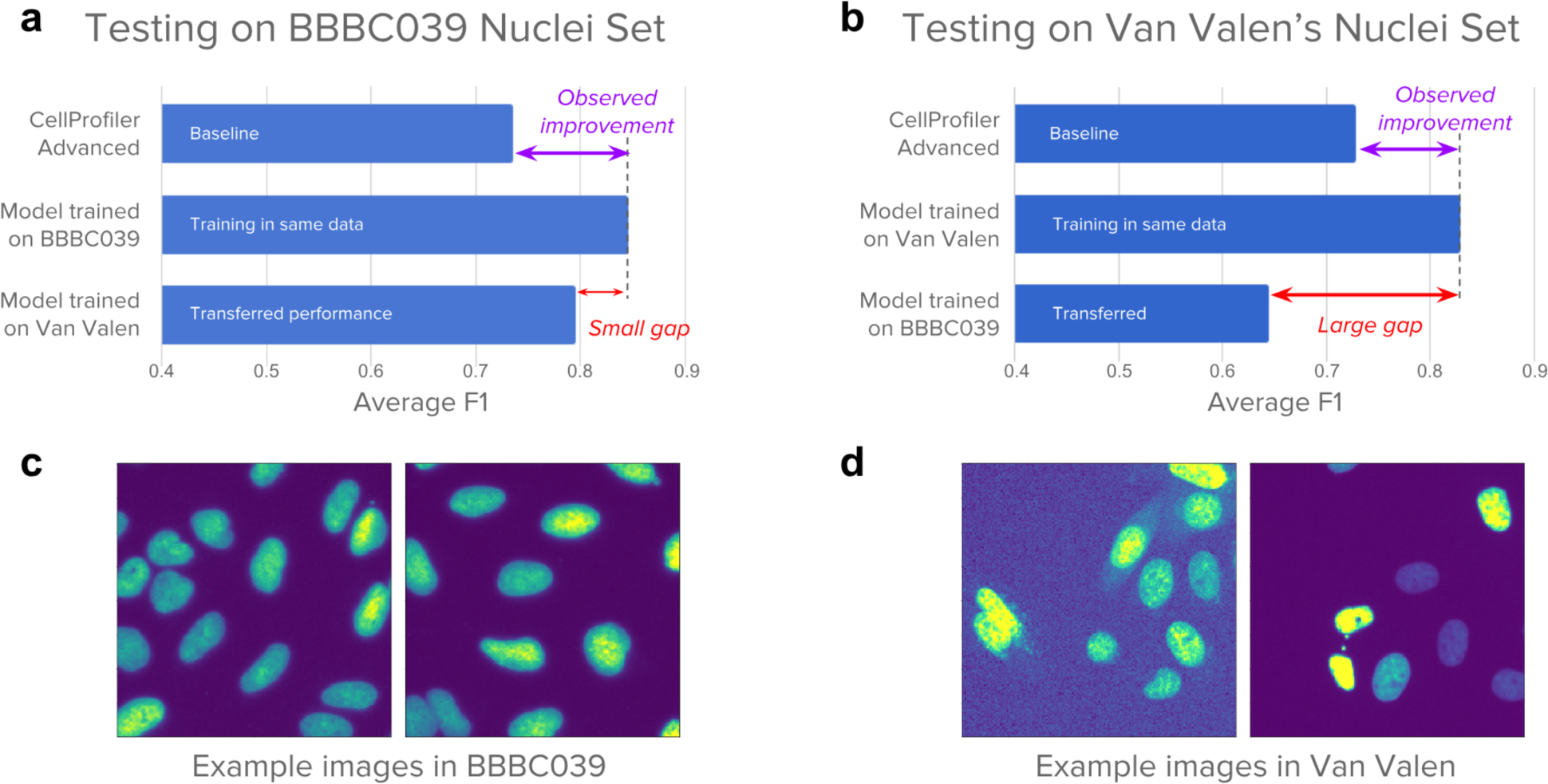
Signal quality is the main challenge when transferring models across experiments: Performance differences when models are trained and evaluated in different experiments. a) Models evaluated on the BBBC039 test set, including a U-Net trained on the same set, another U-Net trained on Van Valen’s set, and a CellProfiler pipeline. The results indicate that transferring the model from one screen to another can bring improved performance. b) Models evaluated in Van Valen’s test set, including CellProfiler baselines adapted to this set, a U-Net trained on the same set, and another U-Net trained on BBBC039. The results illustrate the challenges of dealing with large signal variation. c) Example images from BBBC039 showing homogeneous signal with uniform background. d) Example images from Van Valen’s set illustrating various types of realistic artifacts, such as background noise and high signal variance. Number of training images: 100 in BBBC039 and 9 in Van Valen. Number of test images: 50 in BBBC039 and 3 in Van Valen.

A model trained only on the 9 diverse images of Van Valen’s set generalizes well to test images in BBBC039, improving performance over the baseline (Figure 5a) and reaching comparable performance to the model trained on BBBC039 training images. Note that training a network on images of BBBC039 improves performance with respect to the CellProfiler baseline. The transferred model does not fix all the errors, likely because the number of training examples is limited. Nevertheless, the transferred performance indicates that it is possible to reuse models across experiments to improve segmentation accuracy.

A transfer from the more homogenous BBBC039 set to the more diverse Van Valen set is less successful: a model trained with 100 examples from the BBBC039 set fails to improve on the test set of Van Valen’s images despite the availability of more data (Figure 5b). This demonstrates the challenges of dealing with varying signal quality, which is a frequent concern in high-throughput and high-content screens. The large gap in performance is explained by varying signal conditions (Figure 5c, d): because the model did not observe these variations during training, it fails to correctly segment test images.

The CellProfiler pipelines also confirm the difficulty of handling noisy images. A single pipeline cannot deal with all variations in Van Valen’s test set, requiring the adjustment of advanced settings and the splitting of cases into two different pipelines. In BBBC039, a single pipeline works well due in part to the homogeneity of signal in this collection; the errors are due to challenging phenotypic variations, such as tiny nuclei or clumped objects.

### 4.5 Deep learning needs more computing and annotation time than classical methods

Although we found the performance of deep learning to be favorable in terms of improving segmentation accuracy, we also found that this comes at higher computational cost and annotation time. First, deep learning requires significantly more time to prepare training data with manual annotations (Figure 6a). Second, deep learning needs the researchers to train a model and tune its parameters (Figure 6b), usually with special hardware. Third, when a model has been trained, it is slower to run on new images than classical algorithms (Figure 6c). However, running times can be accelerated using graphic cards, which makes the technique usable in practice.

**Figure 6.**
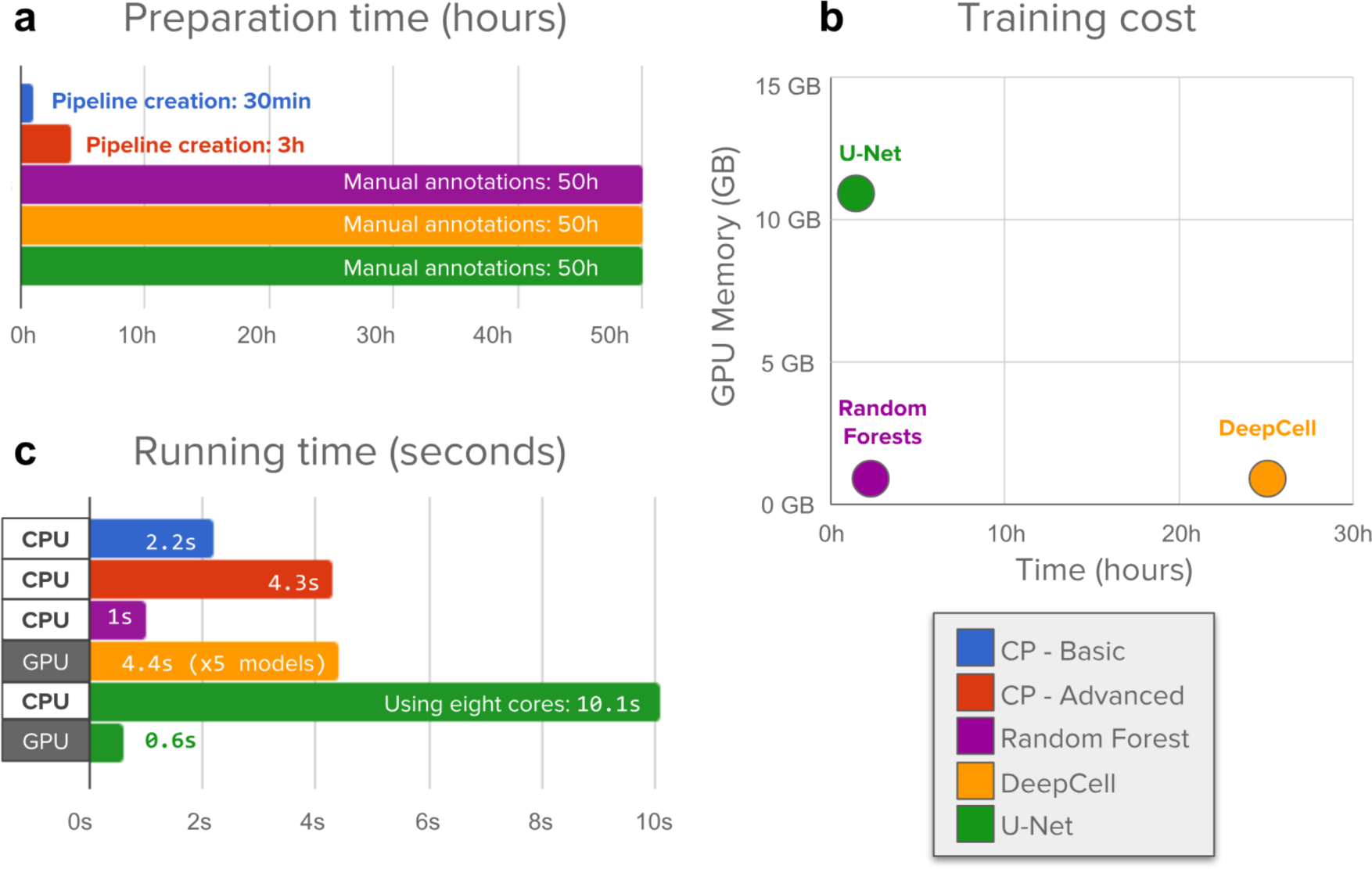
Evaluation of the time needed to create annotations, train, and run segmentation models. a) Preparation time measures hands on, expert time annotating images or creating CellProfiler pipelines. Manually annotating 100 training images with about 11,500 nuclei requires significantly longer times. b) Machine learning models need to be trained while CellProfiler pipelines do not need additional processing. Neural network training was run on a single NVIDIA Titan X GPU. DeepCell trains an ensemble of 5 models, which was used in all evaluations. c) CellProfiler pipelines and Random Forests are run on new images using CPU cores to measure the computational cost of segmenting a single image. Deep learning needs significantly more resources to accomplish the task, but can be accelerated using GPUs, which have thousands of computing cores that allow algorithms to run operations in parallel. This reduces significantly the elapsed time, making it practical and even faster than classical solutions.

We observed that the time invested by experts for annotating images is significantly longer than configuring CellProfiler segmentation pipelines (Figure 6a). We estimate that manually annotating 100 images for training (∼11,500 objects) takes 50 hours of work using an assisted-segmentation tool. In contrast, a basic CellProfiler pipeline can be calibrated in 15 to 30 minutes of interaction with the tool, setting up a configuration that even users without extensive experience nor computational expertise could complete. CellProfiler is very flexible and allows users to add more modules in order to correct certain errors and factor out artifacts, creating an advanced pipeline that can take from 1 to 3 hours.

Training deep learning models takes substantial computing time on GPUs, while CellProfiler pipelines do not need any additional training or post-processing (Figure 6b). In our study, the deep learning models under evaluation are big enough to need a GPU for training, but light enough to be trained in a few hours. In particular, a U-Net can be trained in a single NVIDIA Titan X GPU in just one hour, while DeepCell takes 25 hours for an ensemble of 5 networks (as suggested in the original work (17)). Also, training models may need preliminary experiments to calibrate hyperparameters of the neural network (e.g. learning rate, batch size, epochs), which adds more hands-on time.

Notice the trade-off between memory and training time of deep learning models. DeepCell trains a compact neural network with small memory requirements, which makes it ideal for running experiments efficiently in modest hardware configurations. This design has the additional advantage of allowing strategic pixel sampling for training, resulting in a balanced selection of pixels from the background, boundary, and interior of cells. In contrast, U-Net trains larger neural network models that process all the pixels simultaneously using a fully convolutional approach resulting in high memory requirements and, therefore, needing expensive GPU hardware for training. This can be constraining for laboratories not equipped to run computationally heavy experiments.

When segmenting new images using CPU cores, deep learning models are slower than CellProfiler pipelines (Figure 6c). The computational complexity in terms of space (memory) and time (operations) of a convolutional neural network is proportional to the number of layers, the number of filters, and the size of images. As these architectures get deeper and more complex, they involve more operations to produce the final result, and thus require more computing power. This is in contrast to classical segmentation algorithms whose thresholding and filtering operations have relatively limited computing requirements that scale well with the size of images. Even with 8 times more cores, a U-Net takes 10.1 seconds to segment a single image, which results in about 20 times more computing power requirements than the advanced CellProfiler pipeline. CellProfiler pipelines are run in a single CPU core and take 2.2 and 4.3 seconds for the basic and advanced pipelines respectively.

Using GPU acceleration can significantly speed up the computations of deep learning models, making them very usable and efficient in practice (Figure 6c). Segmenting a single image with a U-Net model takes only 0.6 seconds on a Nvidia Titan X GPU, improving computation times by a factor of 16X. Note that no batching was used for prediction, which can accelerate computation of groups of images even further. In our experiments, a single DeepCell network ran at 4.4 seconds per image with GPU acceleration, which can be further sped up using implementations of dilated convolution and pooling operations available in modern deep learning frameworks. These results show that deep learning models are faster than classical algorithms when using appropriate hardware and efficient implementations.

### 4.6 Better segmentations improve high-content cytometry screens

Accurate nucleus segmentation improves the sensitivity of cytometry screens in real world high-throughput applications. To quantify this effect, we evaluated the performance of the segmentation using the Z’-factor in a high-content experiment. The goal of this experiment is to identify compounds that disrupt normal cell cycle, thus, we measured the DNA content of single cells using the total integrated intensity within each segmented nucleus as a readout.

We selected a subset of 10 compounds screened with high-replicates in the BBBC022 image collection (Figure S3) to measure their effect to the cell cycle compared to DMSO treated wells (negative controls). For each image, we measured the integrated intensity of nuclei and estimated the proportion of cells with 4N DNA content. After controlling for potential batch effects, we observed that none of the compounds yield a sufficiently high Z’-factor that indicates cell cycle disruption. It should be noted that the compounds selected are not known to have effects on the cell cycle - thus, we did not expect them all to yield a detectable phenotype in this assay. However, the measurements were a good choice for our purposes, because we wanted a nucleus-based readout whose baseline quality was not so high so as to leave no room for improvement with better segmentation. For 7 out of the 10 selected compounds we observed improved Z’-factor scores when the segmentation was carried out with a deep learning model, indicating more sensitivity of the screen for detecting interesting compounds.

Other studies have also observed improved assay quality when using deep learning for cell segmentation. For instance, analyzing the structure of tumors in tissues using multiplexed ion beam imaging reveals the spatial organization and response of immune cells in cancer patients (39). These results were obtained with a computational workflow powered by the DeepCell library, demonstrating that deep learning can accurately segment single cells in challenging imaging conditions.

## 5. Discussion

Previous studies of cell segmentation observed minor improvements when comparing deep learning methods vs. classical algorithms using the Jaccard index (17). Also, a recent cell tracking challenge analysis noted that thresholding approaches are still the top solutions for segmenting cells (19); their ranking score is also the Jaccard index. We argue that these evaluation methods do not satisfactorily capture biologically important error modes, making it difficult to appropriately assess cell and nuclei segmentation algorithms. We presented an evaluation framework focused on object level accuracy, which captures biological interpretations more naturally than pixel-level scores (Figure S1).

The objects of interest in biology are full instances of nuclei that experts can identify by eye. Our results show that deep learning strategies improve segmentation accuracy and reduce the number of errors significantly as compared to baselines based on classical image processing and machine learning. However, we show that these methods still make mistakes that experts do not. Being able to quantify these mistakes explicitly will drive future research towards better methods that could match human performance. In our benchmark, deep learning provided improved performance in all the tests that measured accuracy and error rates. Despite requiring significant annotation effort and computational cost, deep learning methods can have a positive impact on the quality and reliability of the measurements extracted from fluorescence images.

### 5.1 Improved accuracy

The proposed evaluation metrics are able to distinguish biologically meaningful errors, helping to better differentiate the performance of segmentation models. The results of our benchmark show that deep learning methods can bring significant advantages for segmenting single objects in new images, compared to manually configured image processing algorithms and classic machine learning. Both DeepCell and U-Net are able to improve the segmentation accuracy (Figure 2 and S1) and reduce the total number of errors (Figure 3 and S2). This shows that even though these models were created for generic cell segmentation and optimized for learning from small datasets, they can be trained to successfully identify nuclei using large sets of images.

The analysis of errors indicates that deep learning can fix most of the segmentation errors observed in classical algorithms, especially merges. One special type of error that represents a challenge for both deep learning models is the segmentation of tiny nuclei (micronuclei and other debris, if of interest in an experiment). In extensive experiments conducted in the DeepCell study (17), the size of the receptive field has been shown to be a key parameter to improve performance, suggesting that crops that fully cover single cells are sufficiently informative for accurate segmentation. Increasing the resolution of images, either during acquisition (40) or with computational methods such as resizing images to make objects look bigger, may help fix these errors. Alternatively, different loss functions adapted to this problem might be designed.

### 5.2 Training data

In our evaluation, the amount of training data was shown to be an important factor to reduce the number of errors. Our results confirm that training a neural network with only a few images is enough to get improved performance relative to non-deep learning baselines. However, in order to improve accuracy and leverage the learning capacity of deep learning models, more data is required. Importantly, a neural network can also be reused across experiments, as long as the training data incorporates variations in morphological phenotypes as well as variations in signal quality and acquisition conditions. We argue that a single deep learning model might be constructed to address all the challenges of nucleus segmentation in fluorescence images if a diverse database of annotated examples were to be collected to incorporate these two critical axes of variation. We advocate for collecting that data collaboratively from different research labs, so everyone will benefit from a shared resource that can be used for training robust neural networks. We have begun such an effort via the 2018 Data Science Bowl https://www.kaggle.com/c/data-science-bowl-2018/.

### 5.3 Computational cost

Deep learning models generally run a higher computational cost. GPUs can be useful in microscopy laboratories for accelerating accurate neural network models; if acquisition or maintenance is prohibitive, cloud computing allows laboratories to run deep learning models using remote computing resources on demand. Adopting these solutions will equip biologists with essential tools for many other image analysis tasks based on artificial intelligence in the future.

## Acknowledgements

We thank Beth Cimini and Minh Doan for their efforts and guidance when annotating the image set used for this research. We also thank Mohammad Rohban and Beth Cimini for fruitful discussions and key insights to design experiments and write the manuscript. We are grateful to David Van Valen for his help while running the DeepCell library and for his constructive suggestions to improve the clarity of the manuscript. Funding was provided by the National Institute of General Medical Sciences of the National Institutes of Health under MIRA award number R35 GM122547 (to AEC).

## Author Contributions

JCC contributed experiments, data analysis, software development, and manuscript writing. JR contributed experiments, data preparation, data analysis and software development. AG contributed experimental design, data preparation and software development. TB contributed experiments, software development and manuscript writing. KWK contributed experiments, data preparation and data analysis. MB contributed experiments and data analysis. MC contributed experiments. CM contributed data preparation and software development. SS contributed experimental design and manuscript writing. FT contributed experimental design and manuscript writing. AEC contributed experimental design, data interpretation and manuscript writing.

## Supplementary Material

### A. Experimental Design

The main dataset in this paper, BBBC039, was split in three subsets, using 50% of the images for training, 25% for validation, and 25% for testing (holdout set). All optimization steps of the deep learning strategies were tuned using the training and validation sets, and the reported results are obtained with final evaluations using the holdout test set (unless otherwise indicated).

We evaluate five segmentation strategies, two based on deep learning, one based on classical machine learning, and two baseline pipelines based on the CellProfiler software. The deep learning strategies are U-Net (16), representing a family of neural networks originally designed for segmentation; and DeepCell (17), representing another family of neural networks for patch-based, pixel-wise classification. Random Forest was selected as the classical machine learning algorithm using the implementation provided by Ilastik (10), which is specialized for segmentation. The baseline segmentation pipelines include an advanced settings mode and a basic settings mode.

With trained deep learning models and calibrated CellProfiler pipelines, we proceeded to segment all images in the test set. To evaluate and compare the estimated masks we use the average F1-score and modes of error.

### B. Post-processing operations

The following are the post-processing steps used to recover single object instances from pixel classification probability maps: 1) max-binarization: the prediction for each pixel is a 3-way probability distribution. These predictions are binarized by assigning 1 to the category with maximum probability and 0 to the rest. 2) Isolation of the interior channel. The other channels only serve the purpose of indicating where the boundaries and background are located, thus they are ignored after determining binary values for each pixel. 3) Use the connected components algorithm (41) to label single objects in the interior channel. After removing the boundaries and background, single nuclei are separated in a binary map. Each of these regions is assigned a different number. 4) Dilation of single objects. The dilation morphological operation (42) is applied to all objects using a square structuring element of 3 × 3 pixels to compensate for the loss of boundaries. This procedure was used for the two evaluated deep learning models: DeepCell and U-Net. No additional boundary refinement algorithms –such as watershed or similar– were applied.

**Supplementary Figure S1.**
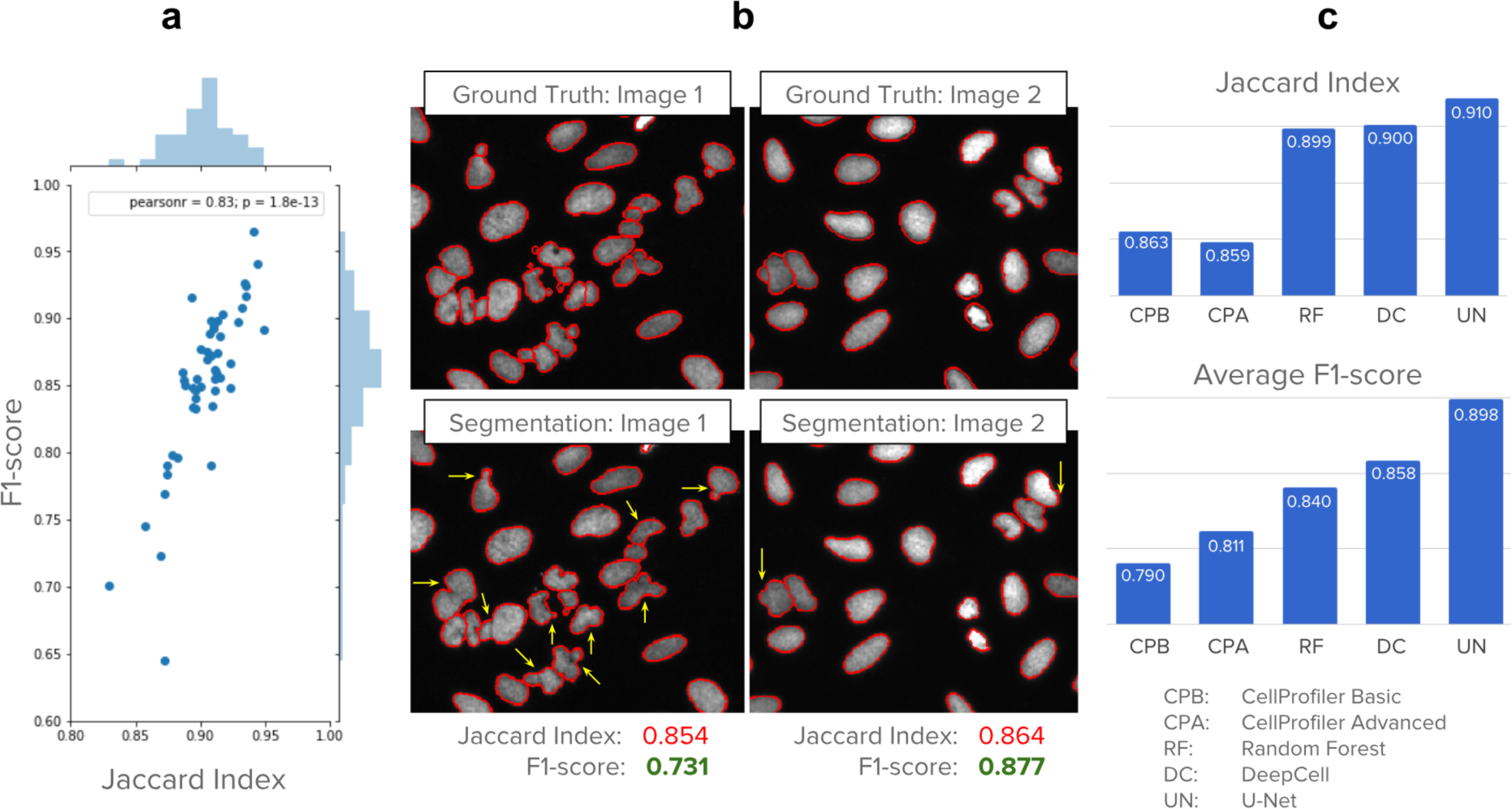
F1-score has better resolution for detecting performance differences than the Jaccard index. In addition, the F1-score is penalized by biologically relevant errors that can be explained in terms of missing, merged and split objects. a) F1-score and Jaccard index are highly correlated measures of segmentation performance, but F1-score displays more resolution to distinguish biologically meaningful errors. Each dot in the plot is an image in the test set segmented with the CellProfiler advanced pipeline. b) Example images that get similar Jaccard indexes but have very different segmentation errors as shown by the yellow arrows. F1-score captures these differences more naturally, which is explained by the number of false negatives resulting from the merges. c) Performance of the evaluated methods on the entire test set. The Jaccard index reports little difference between the two CellProfiler baselines, and also between Random Forest and the two deep learning methods. With average F1-score, the biologically relevant errors are measured more explicitly, and performance differences can be observed more clearly.

**Supplementary Figure S2.**
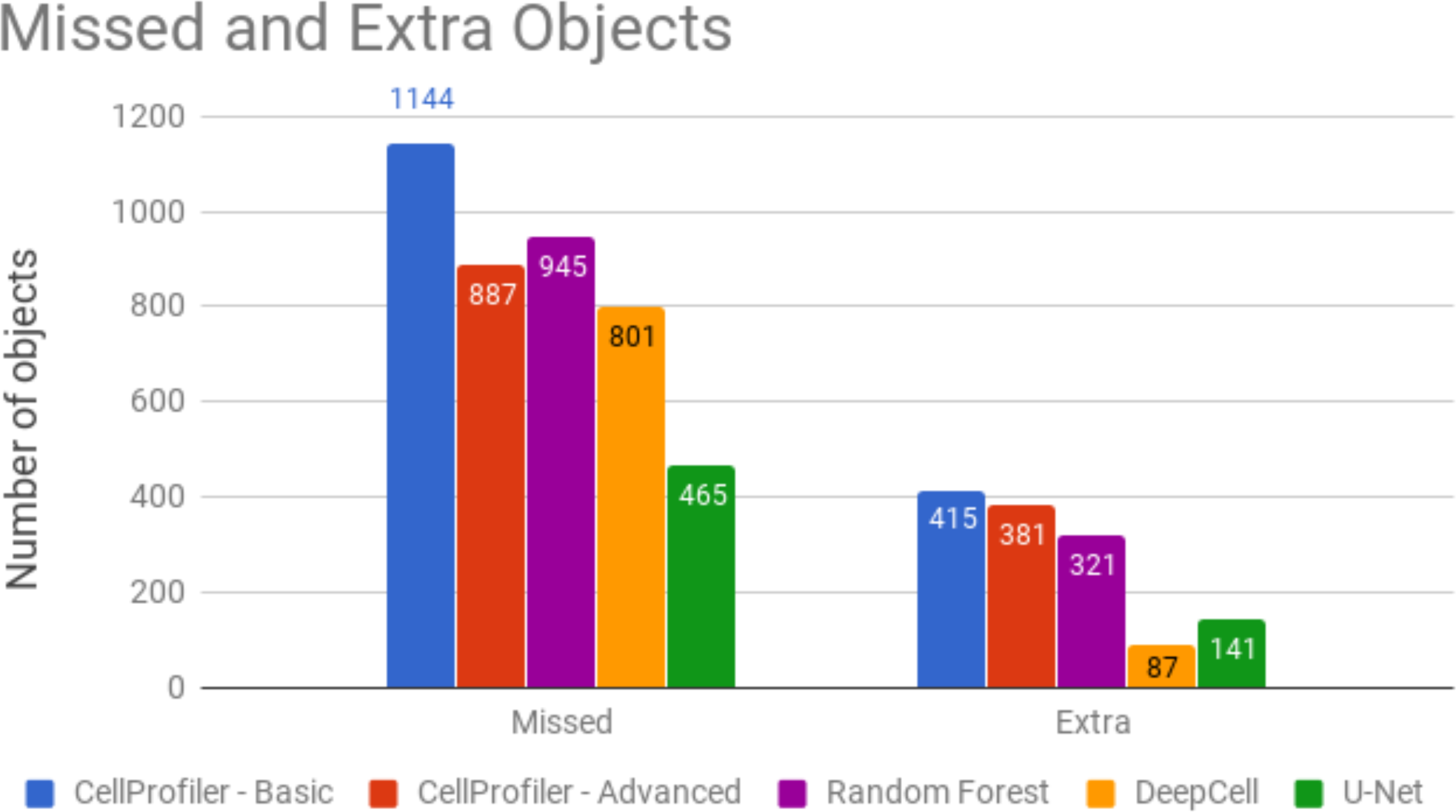
Total number of missed (false negative) and extra (false positive) objects generated by each segmentation model in the test set. The total number of objects in the test set is 5,720. Deep learning algorithms reduce both, missed and extra objects with respect to the baseline. U-Net misses 47% less objects than the CellProfiler advanced pipeline, while Random Forest misses 6% more. Similarly, DeepCell introduces 77% less extra objects, providing cleaner segmentations. Deep learning algorithms display a tradeoff between accurately recovering all relevant objects and not introducing segmentation artifacts (by splits or debris).

**Supplementary Figure S3.**
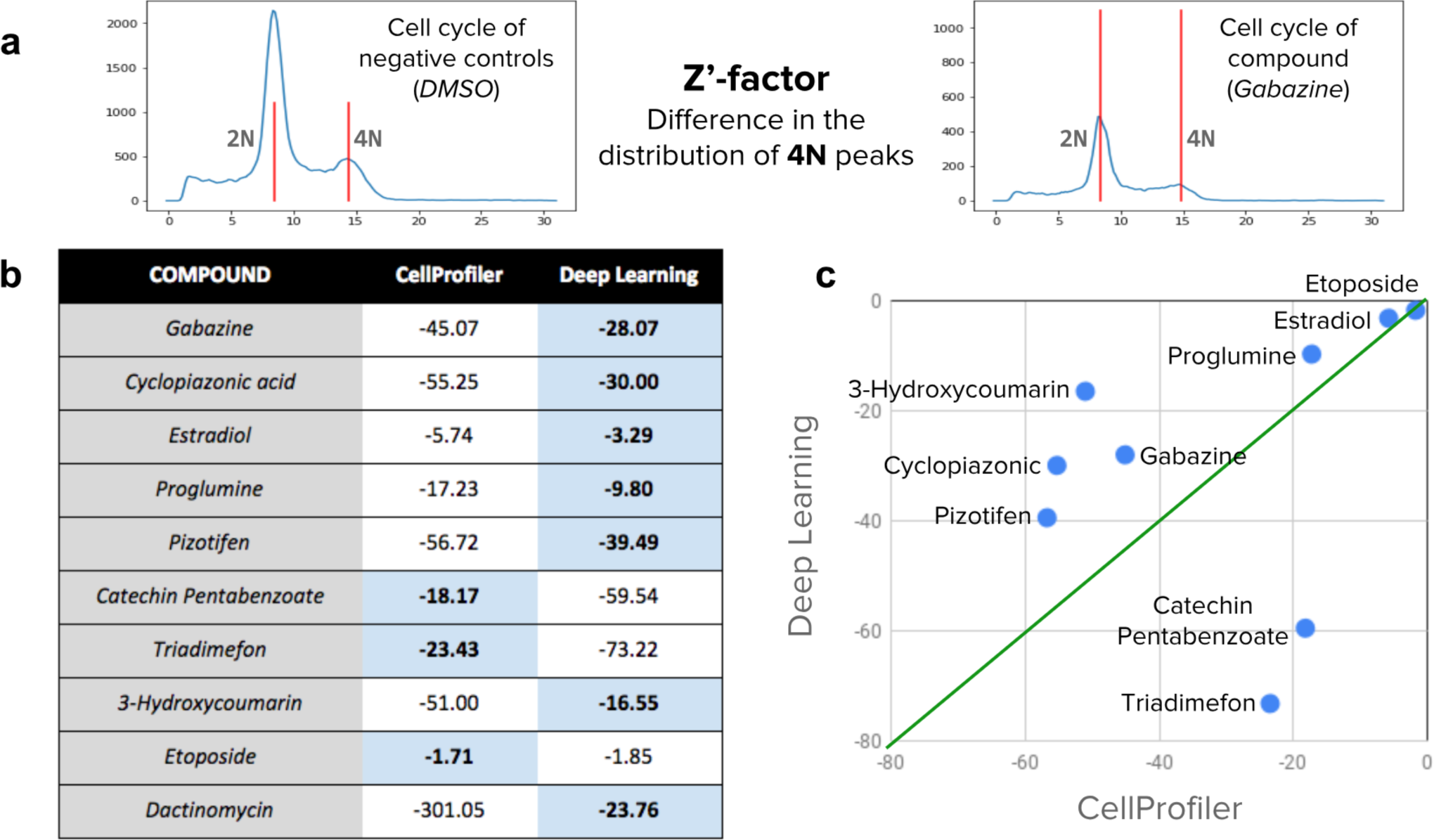
Z’-factor values for a set of 10 compounds selected for evaluation. a) The Z’-factor values indicate whether a compound has an observable cell cycle disruption in a population of cells with respect to negative controls. To compute the Z’-factor, images are first segmented using CellProfiler or a U-Net model. Then, the DNA content is measured within each nucleus using the integrated intensity of pixels. The distribution of DNA content reflects the expected 2N/4N cell cycle. b) Z’-factor values obtained for the two evaluated methods. Higher values are better. c) Scatter plot of the values reported in b. The Y axis presents the result obtained with deep learning and the X axis presents the result with CellProfiler segmentations.

## References

1. Boutros M, Heigwer F, Laufer C. Microscopy-Based High-Content Screening. Cell 2015;163:1314–1325.

2. Mattiazzi Usaj M, Styles EB, Verster AJ, Friesen H, Boone C, Andrews BJ. High-Content Screening for Quantitative Cell Biology. Trends Cell Biol. 2016/8;26:598–611.

3. Caicedo JC, Singh S, Carpenter AE. Applications in image-based profiling of perturbations. Curr. Opin. Biotechnol. 2016;39:134–142.

4. Bougen-Zhukov N, Loh SY, Lee HK, Loo L-H. Large-scale image-based screening and profiling of cellular phenotypes. Cytometry A 2016. Available at: http://dx.doi.org/10.1002/cyto.a.22909.

5. Williams E, Moore J, Li SW, Rustici G, Tarkowska A, Chessel A, Leo S, Antal B, Ferguson RK, Sarkans U, Brazma A, Salas REC, Swedlow JR. The Image Data Resource: A Bioimage Data Integration and Publication Platform. Nat. Methods 2017;14:775–781.

6. Meijering E. Cell Segmentation: 50 Years Down the Road [Life Sciences]. IEEE Signal Process. Mag. 2012;29:140–145.

7. Otsu N. A Threshold Selection Method from Gray-Level Histograms. IEEE Trans. Syst. Man Cybern. 1979;9:62–66.

8. Beucher S, Lantuejoul C. Use of watersheds in contour detection. In: Proceedings of the International Workshop on Image Processing. 1979. Available at: https://ci.nii.ac.jp/naid/10008961959/.

9. Wählby C, Sintorn I-M, Erlandsson F, Borgefors G, Bengtsson E. Combining intensity, edge and shape information for 2D and 3D segmentation of cell nuclei in tissue sections. J. Microsc. 2004;215:67–76.

10. Sommer C, Straehle C, Köthe U, Hamprecht FA. Ilastik: Interactive learning and segmentation toolkit. In: 2011 IEEE International Symposium on Biomedical Imaging: From Nano to Macro. ieeexplore.ieee.org; 2011. p 230–233.

11. Eliceiri KW, Berthold MR, Goldberg IG, Ibáñez L, Manjunath BS, Martone ME, Murphy RF, Peng H, Plant AL, Roysam B, Stuurman N, Stuurmann N, Swedlow JR, Tomancak P, Carpenter AE. Biological imaging software tools. Nat. Methods 2012;9:697–710.

12. Carpenter AE, Jones TR, Lamprecht MR, Clarke C, Kang IH, Friman O, Guertin DA, Chang JH, Lindquist RA, Moffat J, Golland P, Sabatini DM. CellProfiler: image analysis software for identifying and quantifying cell phenotypes. Genome Biol. 2006;7:R100.

13. Schindelin J, Arganda-Carreras I, Frise E, Kaynig V, Longair M, Pietzsch T, Preibisch S, Rueden C, Saalfeld S, Schmid B, Tinevez J-Y, White DJ, Hartenstein V, Eliceiri K, Tomancak P, Cardona A. Fiji: an open-source platform for biological-image analysis. Nat. Methods 2012;9:676–682.

14. LeCun Y, Bengio Y, Hinton G. Deep learning. Nature 2015;521:436–444.

15. Mnih V, Kavukcuoglu K, Silver D, Rusu AA, Veness J, Bellemare MG, Graves A, Riedmiller M, Fidjeland AK, Ostrovski G, Petersen S, Beattie C, Sadik A, Antonoglou I, King H, Kumaran D, Wierstra D, Legg S, Hassabis D. Human-level control through deep reinforcement learning. Nature 2015;518:529–533.

16. Ronneberger O, Fischer P, Brox T. U-net: Convolutional networks for biomedical image segmentation. Med. Image Comput. Comput. Assist. Interv. 2015. Available at: http://link.springer.com/chapter/10.1007/978-3-319-24574-4_28.

17. Van Valen DA, Kudo T, Lane KM, Macklin DN, Quach NT, DeFelice MM, Maayan I, Tanouchi Y, Ashley EA, Covert MW. Deep Learning Automates the Quantitative Analysis of Individual Cells in Live-Cell Imaging Experiments. PLoS Comput. Biol. 2016;12:e1005177.

18. Dima AA, Elliott JT, Filliben JJ, Halter M, Peskin A, Bernal J, Kociolek M, Brady MC, Tang HC, Plant AL. Comparison of segmentation algorithms for fluorescence microscopy images of cells. Cytometry A 2011;79:545–559.

19. Ulman V, Maška M, Magnusson KEG, Ronneberger O, Haubold C, Harder N, Matula P, Matula P, Svoboda D, Radojevic M, Smal I, Rohr K, Jaldén J, Blau HM, Dzyubachyk O, Lelieveldt B, Xiao P, Li Y, Cho S-Y, Dufour AC, Olivo-Marin J-C, Reyes-Aldasoro CC, Solis-Lemus JA, Bensch R, Brox T, Stegmaier J, Mikut R, Wolf S, Hamprecht FA, Esteves T, Quelhas P, Demirel Ö, Malmström L, Jug F, Tomancak P, Meijering E, Muñoz-Barrutia A, Kozubek M, Ortiz-de-Solorzano C. An objective comparison of cell-tracking algorithms. Nat. Methods 2017;14:1141–1152.

20. Rapoport DH, Becker T, Madany Mamlouk A, Schicktanz S, Kruse C. A novel validation algorithm allows for automated cell tracking and the extraction of biologically meaningful parameters. PLoS One 2011;6:e27315.

21. Wienert S, Heim D, Saeger K, Stenzinger A, Beil M, Hufnagl P, Dietel M, Denkert C, Klauschen F. Detection and segmentation of cell nuclei in virtual microscopy images: a minimum-model approach. Sci. Rep. 2012;2:503.

22. Gustafsdottir SM, Ljosa V, Sokolnicki KL, Anthony Wilson J, Walpita D, Kemp MM, Petri Seiler K, Carrel HA, Golub TR, Schreiber SL, Clemons PA, Carpenter AE, Shamji AF. Multiplex cytological profiling assay to measure diverse cellular states. PLoS One 2013;8:e80999.

23. Bray M-A, Singh S, Han H, Davis CT, Borgeson B, Hartland C, Kost-Alimova M, Gustafsdottir SM, Gibson CC, Carpenter AE. Cell Painting, a high-content image-based assay for morphological profiling using multiplexed fluorescent dyes. bioRxiv 2016:049817. Available at: http://biorxiv.org/content/early/2016/04/28/049817. Accessed August 12, 2016.

24. Hughes AJ, Mornin JD, Biswas SK, Beck LE, Bauer DP, Raj A, Bianco S, Gartner ZJ. Quanti.us: a tool for rapid, flexible, crowd-based annotation of images. Nat. Methods 2018;15:587–590.

25. Hariharan B, Arbeláez P, Girshick R, Malik J. Simultaneous Detection and Segmentation. arXiv [cs.CV] 2014. Available at: http://arxiv.org/abs/1407.1808.

26. Shelhamer E, Long J, Darrell T. Fully Convolutional Networks for Semantic Segmentation. arXiv [cs.CV] 2016. Available at: http://arxiv.org/abs/1605.06211.

27. He K, Gkioxari G, Dollár P, Girshick R. Mask R-CNN. In: 2017 IEEE International Conference on Computer Vision (ICCV). 2017. p 2980–2988.

28. Zoph B, Le QV. Neural Architecture Search with Reinforcement Learning. arXiv [cs.LG] 2016. Available at: http://arxiv.org/abs/1611.01578.

29. Bannon D, Moen E, Borba E, Ho A, Camplisson I, Chang B, Osterman E, Graf W, Van Valen D. DeepCell 2.0: Automated cloud deployment of deep learning models for large-scale cellular image analysis. bioRxiv 2018:505032. Available at: https://www.biorxiv.org/content/early/2018/12/22/505032. Accessed January 15, 2019.

30. Nair V, Hinton GE. Rectified linear units improve restricted boltzmann machines. In: Proceedings of the 27th international conference on machine learning (ICML-10). cs.toronto.edu; 2010. p 807–814.

31. Ioffe S, Szegedy C. Batch Normalization: Accelerating Deep Network Training by Reducing Internal Covariate Shift. In: International Conference on Machine Learning. jmlr.org; 2015. p 448–456.

32. Hinton GE, Salakhutdinov RR. Reducing the dimensionality of data with neural networks. Science 2006;313:504–507.

33. McQuin C, Goodman A, Chernyshev V, Kamentsky L, Cimini BA, Karhohs KW, Doan M, Ding L, Rafelski SM, Thirstrup D, Wiegraebe W, Singh S, Becker T, Caicedo JC, Carpenter AE. CellProfiler 3.0: next generation image processing for biology. PLoS Comput. Biol. 2018.

34. Falk T, Mai D, Bensch R, Çiçek Ö, Abdulkadir A, Marrakchi Y, Böhm A, Deubner J, Jäckel Z, Seiwald K, Dovzhenko A, Tietz O, Dal Bosco C, Walsh S, Saltukoglu D, Tay TL, Prinz M, Palme K, Simons M, Diester I, Brox T, Ronneberger O. U-Net: deep learning for cell counting, detection, and morphometry. Nat. Methods 2018. Available at: http://dx.doi.org/10.1038/s41592-018-0261-2.

35. Gudla PR, Nandy K, Collins J, Meaburn KJ, Misteli T, Lockett SJ. A high-throughput system for segmenting nuclei using multiscale techniques. Cytometry A 2008;73:451–466.

36. Everingham M, Van Gool L, Williams CKI, Winn J, Zisserman A. The Pascal Visual Object Classes (VOC) Challenge. Int. J. Comput. Vis. 2010;88:303–338.

37. Lin T-Y, Maire M, Belongie S, Hays J, Perona P, Ramanan D, Dollár P, Zitnick CL. Microsoft COCO: Common Objects in Context. In: Computer Vision – ECCV 2014. Springer International Publishing; 2014. p 740–755.

38. Hoiem D, Chodpathumwan Y, Dai Q. Diagnosing Error in Object Detectors. In: Computer Vision – ECCV 2012. Springer Berlin Heidelberg; 2012. p 340–353.

39. Keren L, Bosse M, Marquez D, Angoshtari R, Jain S, Varma S, Yang S-R, Kurian A, Van Valen D, West R, Bendall SC, Angelo M. A Structured Tumor-Immune Microenvironment in Triple Negative Breast Cancer Revealed by Multiplexed Ion Beam Imaging. Cell 2018;174:1373–1387.e19.

40. Chen F, Tillberg PW, Boyden ES. Optical imaging. Expansion microscopy. Science 2015;347:543–548.

41. Fiorio C, Gustedt J. Two linear time Union-Find strategies for image processing. Theor. Comput. Sci. 1996;154:165–181.

42. Chen S, Haralick RM. Recursive erosion, dilation, opening, and closing transforms. IEEE Trans. Image Process. 1995;4:335–345.

